# Deep mutational scanning of hemagglutinin helps predict evolutionary fates of human H3N2 influenza variants

**DOI:** 10.1101/298364

**Authors:** Juhye M. Lee, John Huddleston, Michael B. Doud, Kathryn A. Hooper, Nicholas C. Wu, Trevor Bedford, Jesse D. Bloom

## Abstract

Human influenza virus rapidly accumulates mutations in its major surface protein hemagglutinin (HA). The evolutionary success of influenza virus lineages depends on how these mutations affect HA’s functionality and antigenicity. Here we experimentally measure the effects on viral growth in cell culture of all single amino-acid mutations to the HA from a recent human H3N2 influenza virus strain. We show that mutations that are measured to be more favorable for viral growth are enriched in evolutionarily successful H3N2 viral lineages relative to mutations that are measured to be less favorable for viral growth. Therefore, despite the well-known caveats about cell-culture measurements of viral fitness, such measurements can still be informative for understanding evolution in nature. We also compare our measurements for H3 HA to similar data previously generated for a distantly related H1 HA, and find substantial differences in which amino acids are preferred at many sites. For instance, the H3 HA has less disparity in mutational tolerance between the head and stalk domains than the H1 HA. Overall, our work suggests that experimental measurements of mutational effects can be leveraged to help understand the evolutionary fates of viral lineages in nature — but only when the measurements are made on a viral strain similar to the ones being studied in nature.

**Significance Statement:** A key goal in the study of influenza virus evolution is to forecast which viral strains will persist and which ones will die out. Here we experimentally measure the effects of all amino-acid mutations to the hemagglutinin protein from a human H3N2 influenza strain on viral growth in cell culture. We show that these measurements have utility for distinguishing among viral strains that do and do not succeed in nature. Overall, our work suggests that new high-throughput experimental approaches may be useful for understanding virus evolution in nature.

---

Seasonal H3N2 influenza virus evolves rapidly, fixing 3 to 4 amino-acid mutations per year in its hemagglutinin (HA) surface protein (1, 2). Many of these mutations contribute to the rapid antigenic drift that necessitates frequent updates to the annual influenza vaccine (3). This evolution is further characterized by competition and turnover among groups of strains called clades bearing different complements of mutations (4–8). Clades vary widely in their evolutionary success, with some dying out soon after emergence and others going on to take over the virus population. Several lines of evidence indicate that successful clades have higher fitness than clades that remain at low frequency (4-6, 9). A key goal in the study of H3N2 evolution is to identify the features that enable certain clades to succeed as others die out.

Two main characteristics distinguish evolutionarily successful clades from their competitors: greater antigenic change, and efficient viral growth and transmission. In principle, experiments could be informative for identifying how mutations affect these features. Most work on influenza evolution to date has utilized experimental data to assess the antigenicity of circulating strains (11–16). However, the non-antigenic effects of mutations also play an important role (5, 7, 9, 17). Specifically, due to influenza virus’s high mutation rate (18–20) and lack of intra-segment recombination (21), deleterious mutations become linked to beneficial ones. The resulting accumulation of deleterious mutations can affect non-antigenic properties central to viral fitness (9). However, there are no large-scale quantitative characterizations of how mutations to H3N2 HA affect viral growth.

It is now possible to use deep mutational scanning (22) to measure the functional effects of all single amino-acid mutations to viral proteins (10, 23-27). However, the only HA for which such large-scale measurements have previously been madewhich the experimental is from the highly lab-adapted A/WSN/1933 (H1N1) strain (10, 23, 24). Here, we measure the effects on viral growth in cell culture of all mutations to the HA of a recent human H3N2 strain. We show that these experimental measurements can help discriminate evolutionarily successful mutations from those found in strains that quickly die out. However, the utility of the experiments for understanding natural evolution depends on the similarity between the experimental and natural strains: measurements made on an H1 HA are not useful for understanding the evolutionary fate of H3 viral strains.

## Results

### Deep mutational scanning of HA from a recent strain of human H3N2 influenza virus

We performed a deep mutational scan to measure the effects of all amino-acid mutations to HA from the A/Perth/16/2009 (H3N2) strain on viral growth in cell culture. This strain was the H3N2 component of the influenza vaccine from 2010-2012 (28, 29). Relative to the consensus sequence for this HA in Genbank, we used a variant with two mutations that enhanced viral growth in cell culture, G78D and T212I (Figure S1 and Dataset S1). The G78D mutation occurs at low frequency in natural H3N2 sequences, and T212 is a site where a mutation to Ala rose to fixation in human influenza in ~2011.

We mutagenized the entire HA coding sequence at the codon level to create mutant plasmid libraries harboring an average of ~1.4 codon mutations per clone (Figure S2). We then generated mutant virus libraries from the mutant plasmids using a helper-virus system that enables efficient generation of complex influenza virus libraries (10) (Figure 1A). These mutant viruses derived all their non-HA genes from the lab-adapted A/WSN/1933 strain. Using WSN/1933 for the non-HA genes reduces biosafety concerns, and also helped increase viral titers. To further increase viral titers, we used MDCK-SIAT1 cells that we had engineered to constitutively express the TMPRSS2 protease, which cleaves the HA precursor to activate it for membrane fusion (30, 31).

**Fig. 1.**
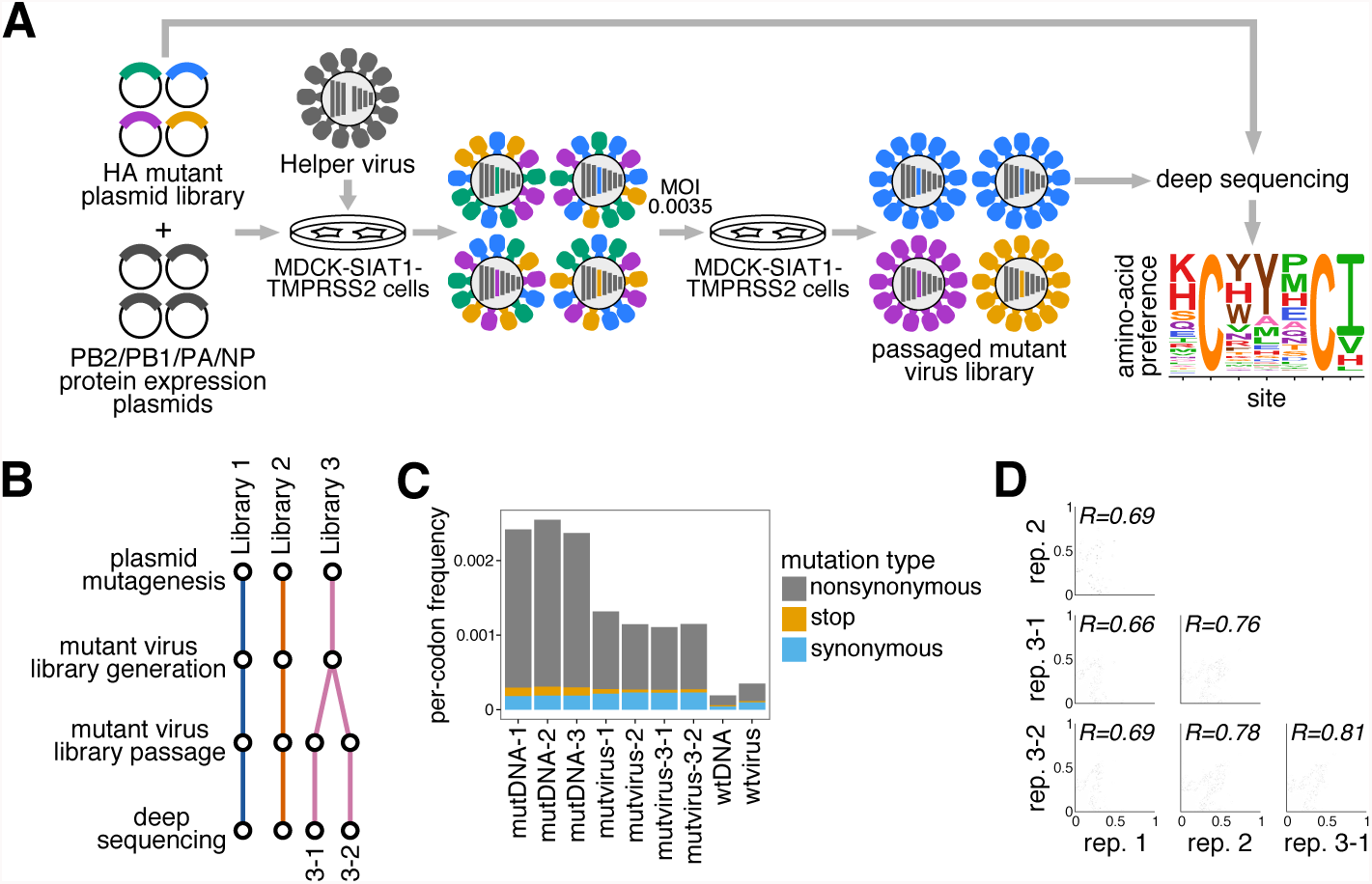
Deep mutational scanning of the Perth/2009 H3 HA. (A) We generated mutant virus libraries using a helper-virus approach (10), and passaged the libraries at low MOI to establish a genotype-phenotype linkage and to select for functional HA variants. Deep sequencing of the variants before and after selection allowed us to estimate each site’s amino-acid preferences. (B) The experiments were performed in full biological triplicate. We also passaged and deep sequenced library 3 in duplicate. (C) Frequencies of nonsynonymous, stop, and synonymous mutations in the mutant plasmid DNA, the passaged mutant viruses, and wildtype DNA and virus controls. (D) The Pearson correlations among the amino-acid preferences estimated in each replicate.

After generating the mutant virus libraries, we passaged them at low MOI in cell culture to create a genotype-phenotype link and select for functional HA variants (Figure 1A). All experiments were completed in full biological triplicate (Figure 1B). We also passaged and deep sequenced library 3 in duplicate (library 3-1 and 3-2) to gauge experimental noise *within* a single biological replicate. As a control to measure sequencing and mutational errors, we used the unmutated HA gene to generate and passage viruses carrying wildtype HA.

Deep sequencing of the initial plasmid mutant libraries and the passaged mutant viruses revealed selection for functional HA mutants. Specifically, stop codons were purged to 20-45% of their initial frequencies after correcting for error rates estimated by sequencing the wildtype controls (Figure 1C). The incomplete purging of stop codons is likely because genetic complementation due to co-infection (32, 33) enabled the persistence of some virions with nonfunctional HAs. We also observed selection against many nonsynonymous mutations (Figure 1C), with their frequencies falling to 30-40% of their initial values after error correction.

We next quantified the reproducibility of our deep mutational scanning across biological and technical replicates. We first used the deep sequencing data for each replicate to estimate the preference of each site in HA for all 20 amino acids as described in (38). Because there are 566 residues in HA, there are 566 x 19 = 10, 754 distinct measurements (the 20 preferences at each site sum to one (38)). The correlations of the amino-acid preferences between pairs of replicates are shown in Figure 1D. The biological replicates were well-correlated, with Pearson’s *R* ranging from 0.69 to 0.78. Replicate 1 exhibited the weakest correlation with other replicates; this replicate also showed the weakest selection against stop and nonsynonymous mutations (Figure 1C), perhaps indicating more experimental noise. The two technical replicates 3-1 and 3-2 were only slightly more correlated than pairs of biological replicates, suggesting that bottlenecking of library diversity during viral passage contributes most of the experimental noise.

### Our measurements are consistent with existing knowledge about HA’s evolution and function

How do the HA amino-acid preferences measured in our experiments relate to the evolution of H3N2 influenza virus in nature? This question can be addressed by evaluating how well an experimentally informed codon substitution model (ExpCM) using our measurements describes H3N2 evolution compared to standard substitution models (34, 39). Table 1 shows that the ExpCM using the across-replicate average of our measurements greatly outperforms conventional substitution models. This result indicates that our experiments authentically capture some of the constraints on HA evolution. The relative rate of nonsynonymous to synonymous substitutions (dN/dS or ω) is <<1 for conventional substitution models (Table 1). However, the relative rate of nonsynonymous to synonymous substitutions after accounting for the amino-acid preferences measured in our experiments (ω for the ExpCM) is close to one (Table 1), indicating that most purifying selection against nonsynonymous substitutions is accounted for in the deep mutational scanning. The ExpCM stringency parameter (34) is 2.47 (Table 1), indicating that natural selection favors the same amino acids as the experiments but with greater stringency. Throughout the rest of this paper, we use the amino-acid preferences rescaled (34, 39) by this stringency parameter. These re-scaled preferences are shown in Figure 2.

**Table 1.**
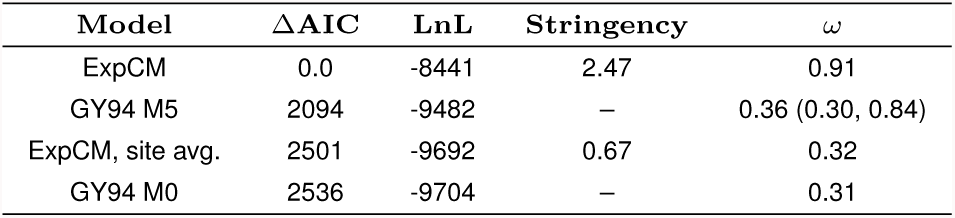
Substitution models informed by the experiments describe HA’s evolution better than traditional models. Maximum likelihood phylogenetic fit to an alignment of human H3N2 HAs using \ (34), ExpCM in which the experimental measurements are averaged across sites (site avg.), and M0 and M5 versions of the Goldman-Yang (GY94) model (35). Models are compared by AIC (36) computed from the log likelihood (LnL) and number of model parameters. The ω parameter is dN/dS for the Goldman-Yang models, and the relative dN/dS after accounting for the measurements for the ExpCM. For the M5 model, we give the mean followed by the shape and rate parameters of the gamma distribution over *ω*.

**Fig. 2.**
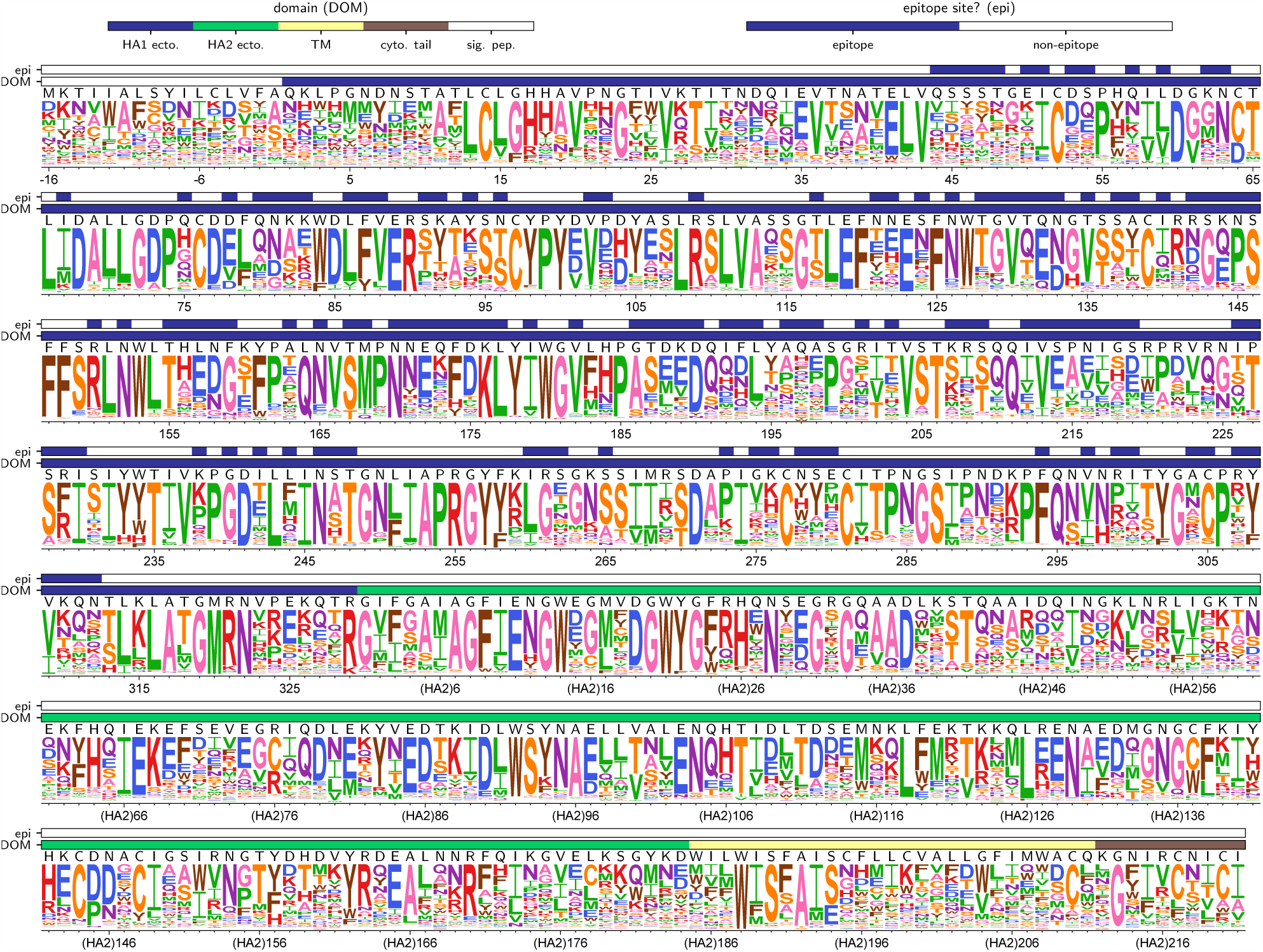
The site-specific amino-acid preferences of the Perth/2009 HA measured in our experiments. The height of each letter is the preference for that amino acid, after taking the average over experimental replicates and re-scaling (34) by the stringency parameter in Table 1. The sites are in H3 numbering. The top overlay bar indicates whether or not a site is in the set of epitope residues delineated in (37). The bottom overlay bar indicates the HA domain (sig. pep. = signal peptide, HA1 ecto. = HA1 ectodomain, HA2 ecto. = HA2 ectodomain, TM = transmembrane domain, cyto. tail. = cytoplasmic tail). The letters directly above each logo stack indicate the wildtype amino acid at that site.

Examination of Figure 2 reveals that the experimentally measured amino-acid preferences generally agree with existing knowledge about HA’s structure and function. For instance, sites that form structurally important disulfide bridges (sites 52 & 277, 64 & 76, 97 & 139, 281 & 305, 14 & 137-HA2, 144-HA2 & 148-HA2) (42) strongly prefer cysteine. At residues involved in receptor binding, there are strong preferences for the amino acids that are known to be involved in binding sialic acid, such as Y98, D190, W153, and S228 (43–46). A positively charged amino acid at site 329 is important for cleavage of the HA0 precursor into the mature form (47), and this site strongly prefers arginine. However, a notable exception occurs at the start codon at position −16, which does not show a strong preference for methionine. This codon is part of the signal peptide and is cleaved from the mature HA protein. One possible reason that our experiments do not show a strong preference for methionine at this site could be alternative translation-initiation at a downstream or upstream start site, as has been described for other HAs (48).

### There is less difference in mutational tolerance between the HA head and stalk domains for H3 than for H1

Our experiments measure which amino acids are tolerated at each HA site under selection for viral growth. We can therefore use our experimentally measured amino-acid preferences to calculate the inherent mutational tolerance of each site, which we quantify as the Shannon entropy of the re-scaled preferences. In prior mutational studies of H1 HAs, the stalk domain was found to be substantially less mutationally tolerant than the globular head (10, 23, 24, 49).

We performed a similar analysis using our new data for the Perth/2009 H3 HA. Surprisingly, the head domain of the H3 HA is *not* more mutationally tolerant than the stalk domain (Figure 3). Specifically, whereas solvent-exposed residues in the head domain are substantially more mutationally tolerant than those in the stalk domain for the WSN/1933 H1 HA, the trend is actually reversed for the Perth/2009 H3 HA (Figure 3B). This difference between the relative mutational tolerances of the H1 and H3 HAs is robust to the cutoff used to define surface residues (Figure S5). For instance, for the H3 HA, the short helix A in the stalk domain is as mutationally tolerant as many surface-exposed residues in the head domain—something that is not the case for the H1 HA. Helix A forms part of the epitope of many broadly neutralizing anti-stalk antibodies (50–52), and interestingly, there are more reports of readily selecting escape mutants from such anti-stalk antibodies in H3 (52–55) than in H1 (56–58) HAs.

**Fig. 3.**
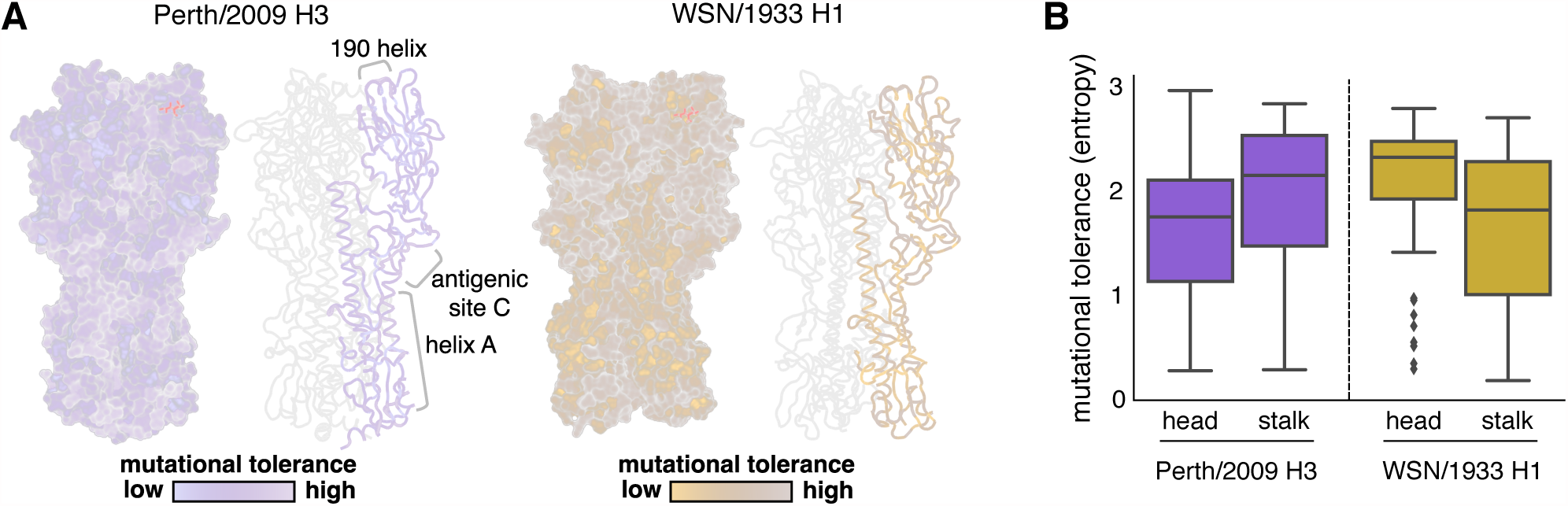
Mutational tolerance of each site in H3 and H1 HAs. (A) Mutational tolerance as measured in the current study is mapped onto the structure of the H3 trimer (PDB 4O5N; (40)). Mutational tolerance of the WSN/1933 H1 HA as measured in (10) is mapped onto the structure of the H1 trimer (PDB 1RVX; (41)). Different color scales are used because measurements are comparable among sites within the same HA, but not necessarily across HAs. Both trimers are shown in the same orientation. For each HA, the structure at left shows a surface representation of the full trimer, while the structure at right side shows a ribbon representation of just one monomer. The sialic acid receptor is shown in red sticks. (B) The mutational tolerance of solvent-exposed residues in the head and stalk domains of the Perth/2009 H3 HA (purple) and WSN/1933 H1 HA (gold). Residues falling in between the two cysteines at sites 52 and 277 were defined as belonging to the head domain, while all other residues were defined as the stalk domain. A residue was classified as solvent exposed if its relative solvent accessibility was ≤ 0.2; Figure S5 shows that the results are robust to the choice of solvent accessibility cutoff. Note that the mutational tolerance values are not comparable between the two HAs, but are comparable between domains of the same HA.

We also see high mutational tolerance in many of the known antigenic regions of H3 HA (59). For instance, antigenic region B is an immunodominant area, and many recent major antigenic drift mutations have occurred in this region (14, 15, 60). We find that the most distal portion of the globular head near the 190-helix, which is part of antigenic region B, is highly tolerant of mutations (Figure 3A). Antigenic region C is also notably mutationally tolerant.

Many residues inside HA’s receptor binding pocket are known to be highly functionally constrained (44, 61), and our data indicates that these sites are relatively mutationally intolerant in both H3 and H1 HAs (Figure 3A). In contrast, the residues surrounding the receptor binding pocket are fairly mutationally tolerant, which may contribute to the rapidity of influenza‘s antigenic evolution, since mutations at these sites can have large effects on antigenicity (14, 59).

### Our measurements can help distinguish between mutations that reach low and high frequencies in nature

Mutations occurring in the H3N2 virus population experience widely varying evolutionary fates (Figure 4). Some mutations appear, spread and fix in the population, while others briefly circulate before disappearing. We take the maximum frequency reached by a mutation as a coarse indicator of its effect on fitness, since favorable mutations generally reach higher frequency than unfavorable ones (62). Here, we follow the population genetic definition of *mutation* and track the outcome of each individual mutation event, e.g. although R142G occurs multiple times on the phylogeny we track each of these mutations occurring on different backgrounds separately. As such, each mutation is shown as a separate circle on a separate branch in Figure 4. However, because multiple mutations on the same phylogeny branch cannot be disentangled, when multiple mutations occurred on a single branch, we assigned a single mutational effect based on the sum of effects of each mutation.

**Fig. 4.**
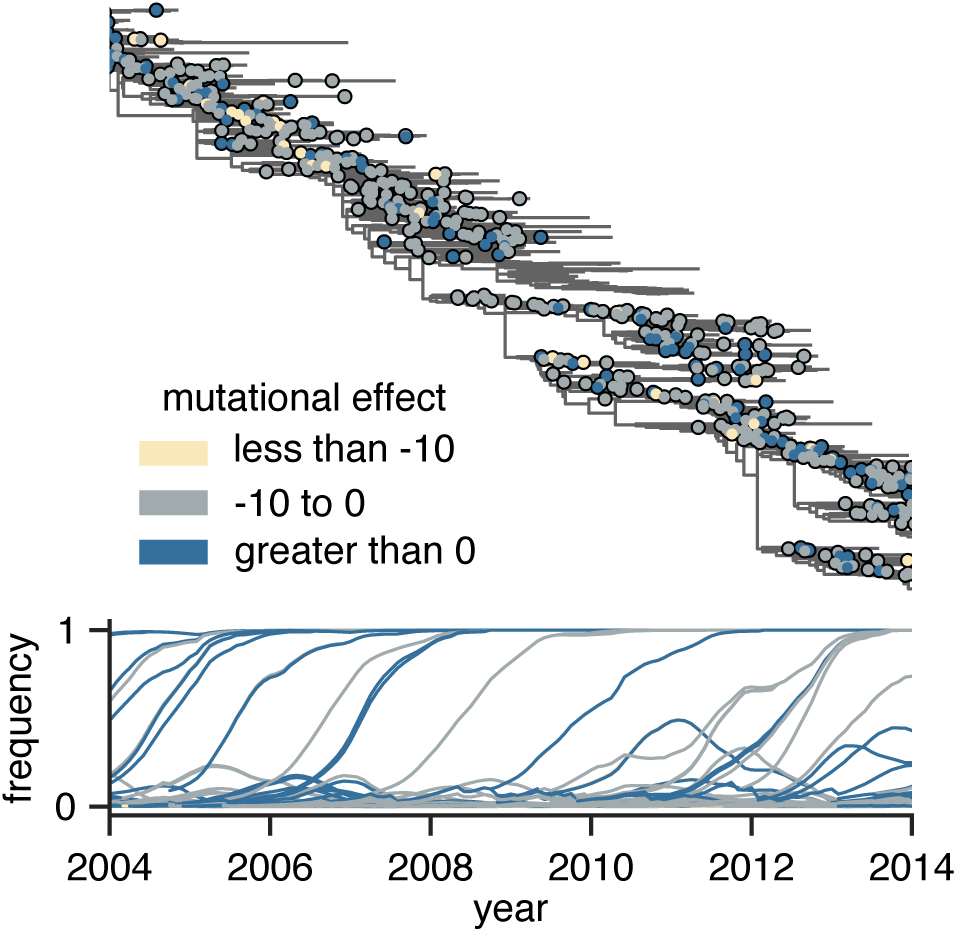
Frequency trajectories of individual mutations and their relation to the experimentally measured effects of these mutations. The top panel shows the subset of the full H3N2 HA tree in Figure S4 from 2004 to 2014. Circles indicate individual amino-acid mutations, and are colored according to the mutational effect measured in our deep mutational scanning (negative values indicate mutations measured to be deleterious to viral growth). The bottom panel shows the frequency trajectory of each mutation, with trajectories colored according to the mutational effects as in the top panel. It is clear that most mutations that reach high frequency are measured to be relatively favorable in our experiments.

After annotating mutations and their frequencies on the phylogeny in this way (Figure 4), it is visually obvious that there are relatively few circulating mutations that we measure to be strongly deleterious—and that such deleterious mutations rarely reach high frequency when they do occur. Mutations that reach high frequencies generally have neutral or beneficial effects according to our experimental measurements.

We next sought to quantify the correlation between a mutation’s experimentally measured effect and the maximum frequency it attained during natural evoluton. To minimize effects related to the genetic background of the strain used in the experiment, we excluded mutations closely related to the experimental strain itself and partitioned the remaining mutations into 1,022 mutations pre-dating and 299 mutations post-dating the Perth/2009 strain (Figure S4). We additionally excluded mutations from the post-Perth partition that were sampled in 2014 or after that have not had enough time for their evolutionary fates to be fully resolved. We used these pre-Perth and post-Perth partitions to test the utility of our measurements for both post-hoc and prospective analyses, respectively. We quantified the relationship between mutational effects and maximum mutation frequencies in the H3N2 phylogeny via Spearman rank correlation (Figure 5A). In both pre-Perth and post-Perth time periods, we found a modest, but statistically significant relationship between mutational effect and maximum mutation frequency (pre-Perth *ρ* = 0.17, post-Perth *ρ =* 0.14). The similar effect sizes for both partitions shows that our experimental measurements can help explain the evolutionary fates of mutations in strains that post-date the experimentally studied strain, as well as to retrospectively analyze mutations that precede the experimental strain.

**Fig. 5.**
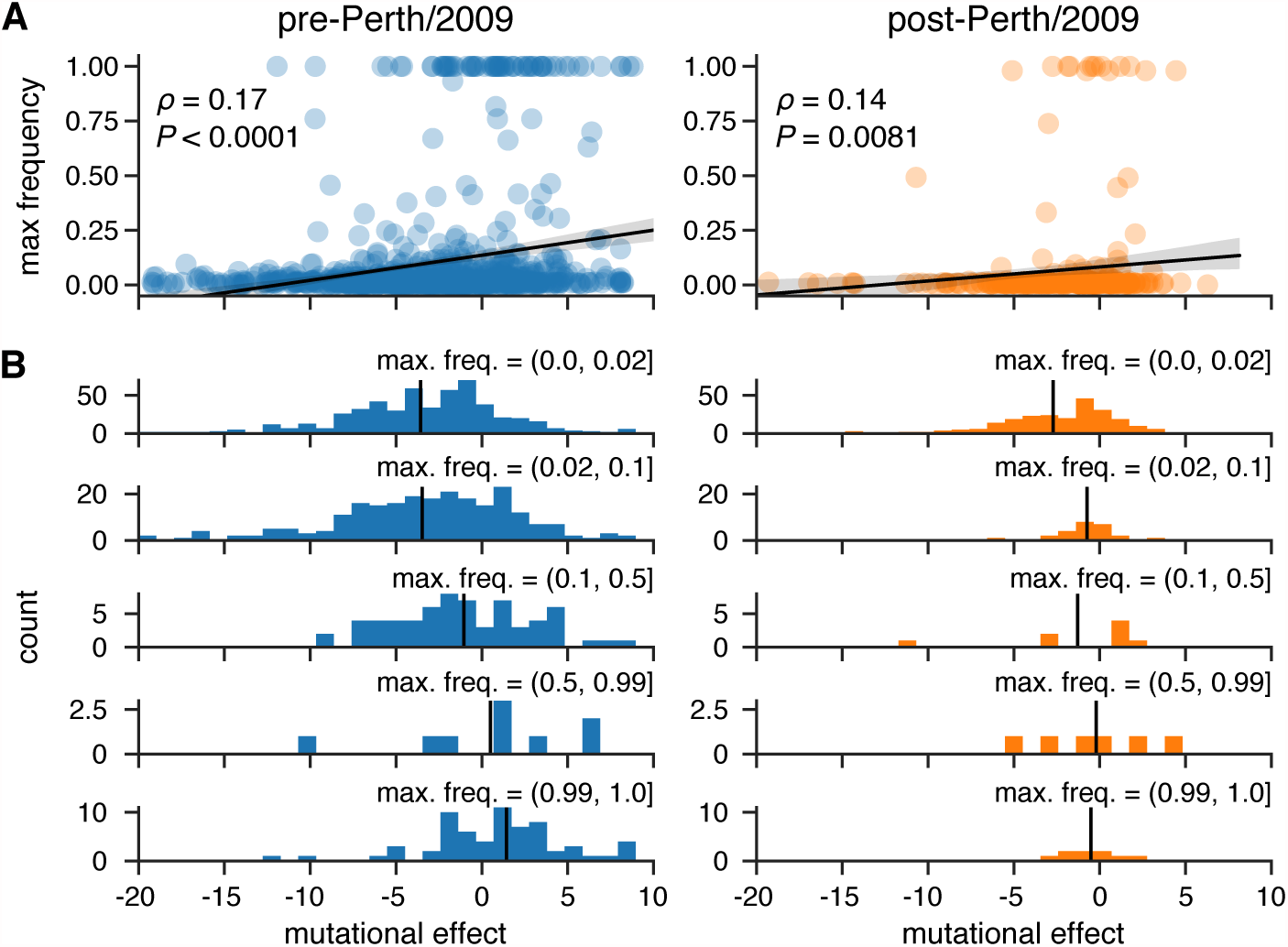
Experimental measurements are informative about the evolutionary fate of viral mutations. (A) Correlation between the effects of mutations as measured in our deep mutational scanning of the Perth/2009 HA and the maximum frequency reached by these mutations in nature. The plots show Spearman *p* and an empir-ical *P*-value representing the proportion of 10,000 permutations of the experimental measurements for which the permuted *p* was greater than or equal to the observed *p*. (B) The distribution of mutational effects partitioned by maximum mutation frequency. The vertical black line shows the mean mutation effect for each category. The analysis is performed separately for the pre- and post-Perth/2009 partitions of the tree (see Figure S4).

The trends in Figure 5A are most strongly driven by the behavior of substantially deleterious mutations. We investigated this further by partitioning mutations into those that reach low, medium and high frequencies, and those that fix in the population (Figure 5B). The mutations that reach higher frequency have a more favorable mean effect. Mutations measured to be substantially deleterious almost never reach high frequency. Overall, these results demonstrate that measurements of how mutations affect viral growth in cell culture are informative for understanding the fates of these mutations in nature: in particular, if a mutation is measurably deleterious to viral growth, that mutation is unlikely to prosper in nature.

### Measurements made on an H1 HA are not useful for understanding the evolution of H3 influenza

To determine how broadly experimental measurements can be generalized across HAs, we repeated the foregoing analysis of H3N2 mutation frequencies, but using mutational effects measured in our prior deep mutational scanning of the WSN/1933 H1 HA, which is highly diverged from the Perth/2009 H3 HA (the two HAs only have 42% protein sequence identity). Figure 6 shows that there is no correlation between the H1 experimental measurements and maximum frequency that mutations reach during H3N2 viral evolution. Therefore, the utility of an experiment for understanding natural evolution degrades as the experimental sequence becomes more diverged from the natural sequences that are being studied.

**Fig. 6.**
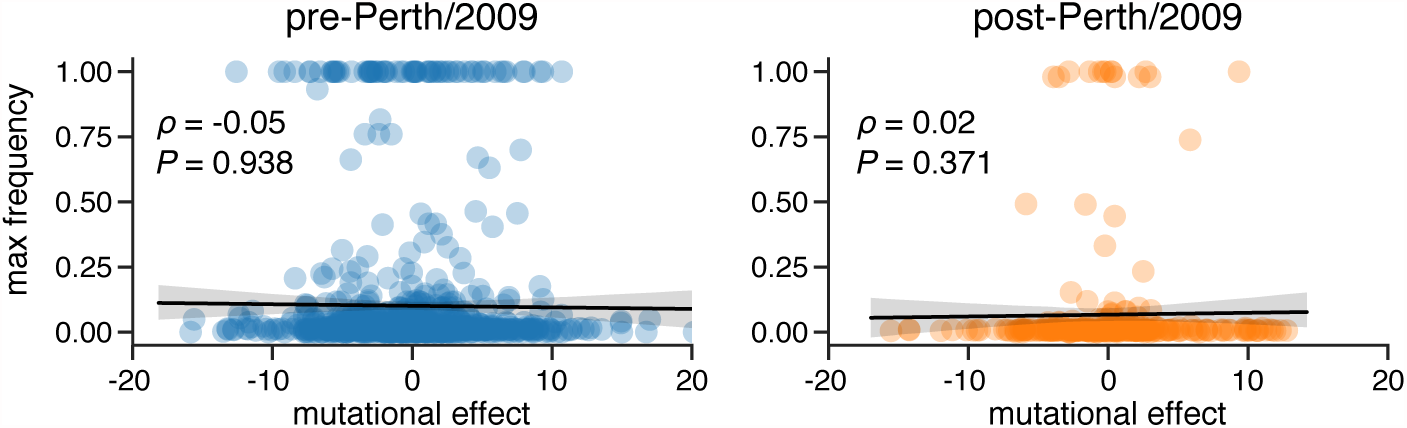
Experimental measurements on an H1 HA are not informative about the evolutionary fate of H3N2 mutations. This figure repeats the analysis of the H3N2 mutation frequencies in Figure 5A, but uses the deep mutational scanning data for an H1 HA as measured in (10), i.e. Figure S6 shows the histograms comparable to those in Figure 5B. The empirical *P*-value represents the results of 1,000 permutations.

### There are large differences between H3 and H1 HAs in the amino-acid preferences of many sites

An obvious hypothesis for why the H1 deep mutational scanning is not useful for understanding the evolution of H3N2 influenza viruses is that the effect of the same mutation is often different between these two HA subtypes. To determine if this is the case, we examined how much the amino-acid preferences of homologous sites have shifted between H3 and H1 HAs. Prior experiments have found only modest shifts in amino-acid preferences between two variants of influenza nucleoprotein with 94% amino-acid identity (63) and variants of HIV envelope (Env) with 86% amino-acid identity (27). However, the H1 and H3 HAs are far more diverged, with only 42% amino-acid identity (Figure 7A). One simple way to investigate the extent of shifts in amino-acid preferences is to correlate measurements from independent deep mutational scanning replicates on H1 and H3 HAs. Figure 7B shows that replicate measurements on the same HA variant are more correlated than those on different HA variants.

**Fig. 7.**
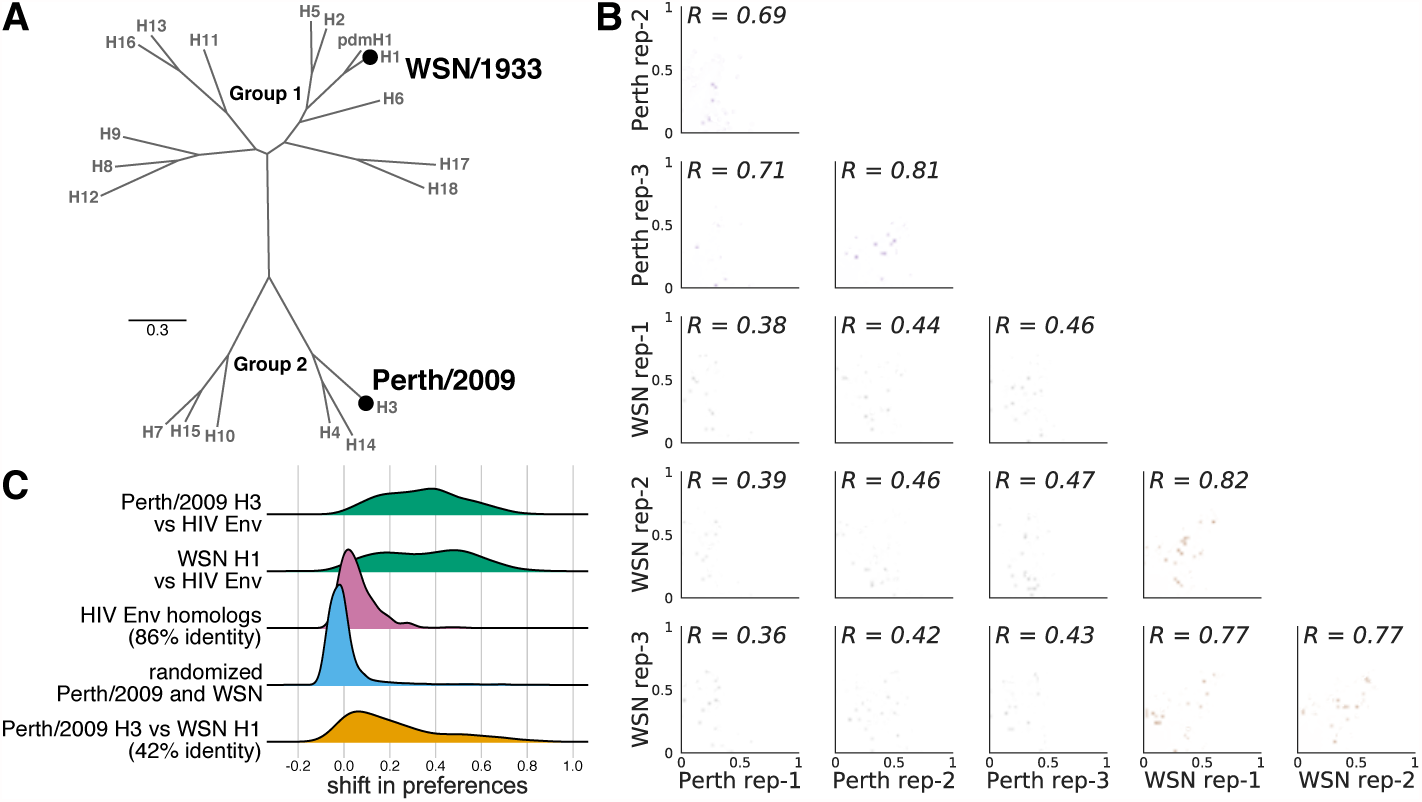
There are large shifts in the effects of mutations between H1 and H3 HAs. (A) Phylogenetic tree of HA subtypes, with the WSN/1933 H1 and Perth/2009 H3 HAs labeled. (B) All pairwise correlations of the amino-acid preferences measured in the three individual deep mu-tational scanning replicates in the current study and the three replicates in prior deep mutational scanning of the H1 HA (10). Comparisons between H3 replicates are in purple, those between H1 replicates are in brown, and those across H1 and H3 replicates are in gray.*R* indicates the Pearson correlation coefficient. (C) We calculated the shift in amino-acid preferences at each site between H3 and H1 HAs using the method in (27), and plotted the distribution of shifts for all sites. The shifts between H3 and H1 (yellow) are much larger than the null distribution (blue) expected if all differences are due to experimental noise. The shifts are also much larger than those previously observed between two variants of HIV Env that share 86% amino-acid identity (pink). However, the shifts between H3 and H1 are less than the differences between HA and HIV Env (green).

To more rigorously quantify shifts in amino-acid preferences after correcting for experimental noise, we used the statistical approach in (27, 63). Figure 7C shows the distribution of shifts in amino-acid preferences between H3 and H1 HAs after correcting for experimental noise. Although some sites have small shifts near zero, many sites have large shifts. These shifts between H3 and H1 are much larger than expected from the null distribution that would be observed purely from experimental noise. They are also much larger than the shifts previously observed between two HIV Envs with 86% aminoacid identity (27). However, the typical shift between H3 and H1 is still smaller than that observed when comparing HA to the non-homologous HIV Env protein. Therefore, there are very substantial shifts in mutational effects between highly diverged HA homologs, although the effects of mutations remain more similar than for non-homologous proteins.

### Properties associated with the shifts in amino-acid preferences between H3 and H1 HAs

What features distinguish the sites with shifted amino-acid preferences between H3 and H1 HAs? The sites of large shifts do not obviously localize to one specific region of HA’s structure (Figure 8A). However, at the domain level, sites in HA’s stalk tend to have smaller shifts than sites in HA’s globular head (Figure 8B). The HA stalk domain is also more conserved in sequence (64), suggesting that conservation of amino-acid sequence is correlated with conservation of amino-acid preferences. Consistent with this idea, sites that are absolutely conserved across all 18 HA subtypes are significantly less shifted than sites that are variable across HA subtypes (Figure 8B). Presumably these sites are under consistent functional constraint across all HAs.

Despite their high sequence divergence, H1 and H3 adopt very similar protein folds (65, 66). However, there are differences in the rotation and upward translation of the globular head subdomains relative to the central stalk domain among different HA subtypes (65, 66). Previous work has defined clades of structurally related HA subtypes (65, 66). One such clade includes H1, H2, H5, and H6, whereas another clade includes H3, H4, and H14 HAs (Figure 7A). Sites that are conserved at different amino-acid identities in these two clades tend to have exceptionally large shifts in amino acid preferences (Figure 8B). The clade containing H1 has an upward translation of the globular head relative to the clade containing H3. This structural shift has been attributed largely to the interaction between sites 107 and 75(HA2) (65, 66). Specifically, the clade containing H1 has a taller turn in the interhelical loop connecting helix A and helix B in the stalk domain, and this tall turn is stabilized by a hydrogen bond between Glu-107 and Lys-75(HA2) (Figure 8C). In deep mutational scanning of the H1 HA, site 107 has a high preference for Glu and 75(HA2) strongly prefers positively charged Lys and Arg. In contrast, the interhelical loop in H3 HA makes a sharper and shorter turn which is facilitated by a Gly at 75(HA2). In the deep mutational scanning of the Perth/2009 H3 HA, site 75(HA2) prefers Gly and to a lesser extent Val, while site 107 is fairly tolerant of mutations. Therefore, some of the shifts in HA amino-acid preferences can be rationalized in terms of changes in HA structure.

**Fig. 8.**
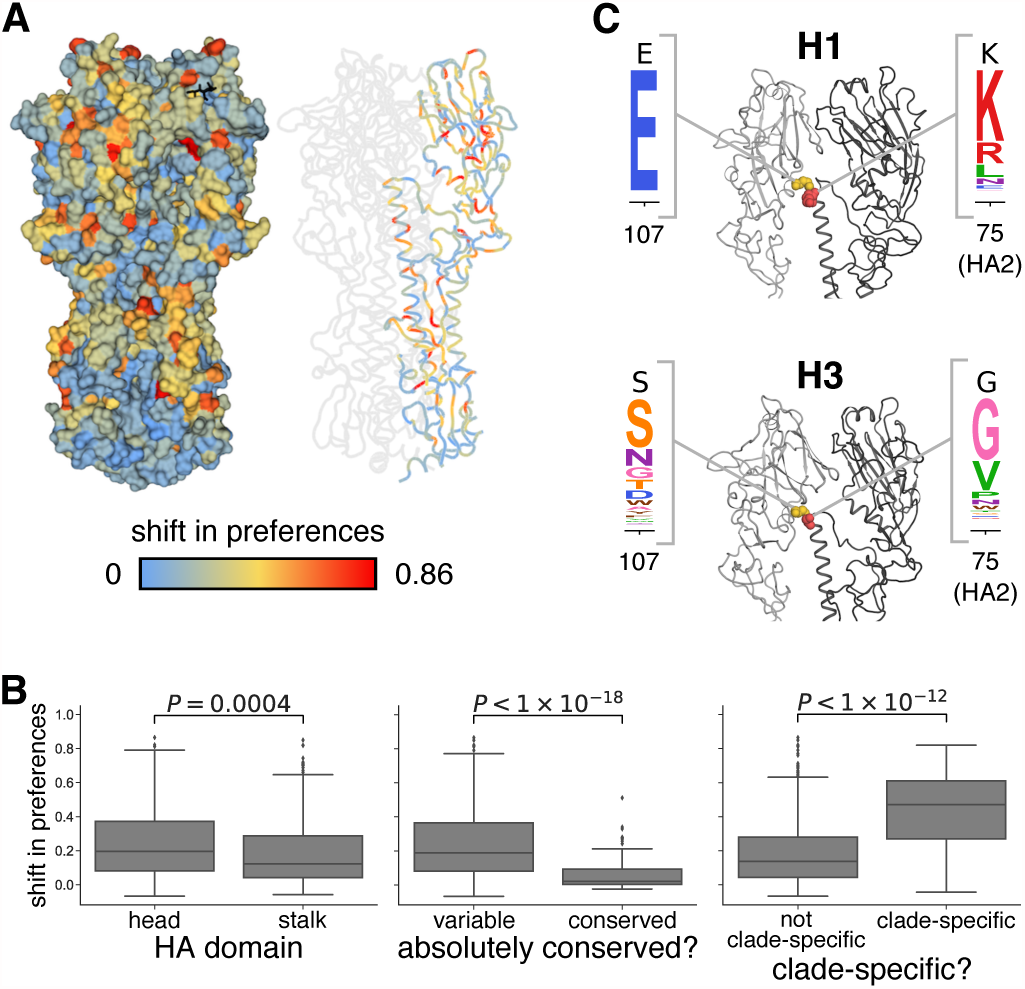
Sites with strongly shifted amino-acid preferences between H3 and H1 HAs. (A) The shift in amino-acid preferences between the H3 and H1 HA at each site as calculated in Figure 7C is mapped onto the structure of the H3 HA. (B) Amino-acid preferences of sites in the stalk domain are less shifted than those in the head domain. Sites absolutely conserved in all 18 HA subtypes are less shifted than other sites. Sites with one amino-acid identity in the clade containing H1, H2, H5, and H6 and another identity in the clade containing H3, H4, and H14 are more shifted than other sites. (C) Sites 107 and 75(HA2) help determine the different orientation of the globular head domain in H1 versus H3 HAs. These sites are shown in spheres on the structure of H1 and H3 and colored as in panel (A), and the experimentally measured amino-acid preferences in the H1 and H3 HAs are shown. One monomer is in dark gray, while the HA1 domain of the neighboring monomer is in lighter gray.

## Discussion

We have measured the effects of all possible single aminoacid mutations to the Perth/2009 H3 HA on viral growth in cell culture and demonstrated that these measurements have some value for understanding the evolutionary fate of these mutations in nature. Specifically, mutations measured to be more beneficial for viral growth tend to reach higher frequencies in nature than mutations measured to be more deleterious for viral growth. The fact that our experiments can help identify evolutionary successful mutations suggests that they might inform evolutionary forecasting. In their landmark paper introducing predictive viral fitness models that accounted for both antigenic and non-antigenic mutations, Łuksza and Lässig (9) noted that the models could in principle be improved by integrating “diverse genotypic and phenotypic data” that more realistically represented the effects of specific mutations. Our work suggests that deep mutational scanning may be able to provide such data.

It is important to emphasize that measurements of viral growth in cell culture do *not* represent true fitness in nature.

Indeed, a vast amount of work in virology has chronicled the ways in which experiments can select for lab artifacts or fail to capture important pressures that are relevant in nature (67-70). Mutations in viral genes other than HA are also important in determining strain success (71, 72). Given these caveats, it might seem surprising that measuring viral growth in cell culture can be informative about the success of viral strains in nature. Yet, prior to our work, there were no comprehensive studies of the functional effects of mutations to H3 HA on any property that even resembled viral fitness in nature, and modeling work has either omitted the non-antigenic effects of mutations (11–13) or assumed that all non-epitope mutations had equivalent deleterious effects (9). The strength of our measurements are not that they perfectly capture fitness in nature, but that they are systematic and quantitative—and so represent an improvement over no information at all. We suspect that performing similar experiments using more realistic and complex selections (e.g., ferrets or primary human airway cultures) might further improve their utility.

We measured the effects of all single amino-acid mutations to a specific HA, and then generalized these measurements to other H3N2 HAs from a 50-year timespan. These generalizations will only be valid to the extent that the effects of mutations are conserved during HA’s evolution. Extensive prior work has shown that epistasis can shift the effects of mutations as proteins evolve (73–79). Our work suggests that measurements on a HA from single human H3N2 viral strain can be usefully generalized to at least some extent across the entire evolutionary history of human H3N2 HA. On the other hand, when we compared our measurements for an H3 HA to prior measurements on H1 HA, we found substantial shifts at many sites—much greater than those observed in prior protein-wide comparisons of more closely related homologs (27, 63). Further investigation of how mutational effects shift as proteins diverge will be important for determining how broadly any given experiment can be generalized when attempting to make evolutionary forecasts.

Our work did not characterize the antigenic effects of mutations, which also play an important role in determining strain success in nature (13, 14). However, our basic selection and deep-sequencing approach can be harnessed to completely map how mutations affect antibody recognition (57, 80). But so far, experiments using this approach have not examined antibodies or sera that are relevant to driving the evolution of H3N2 influenza (57, 80), or have used relevant sera but examined a non-comprehensive set of mutations (16). Future experiments that completely map how HA mutations affect recognition by human sera seem likely to be especially fruitful for informing viral forecasting.

## Materials and Methods

### Data and computer code

Deep sequencing data are available from the Sequence Read Archive under BioSample accessions SAMN08102609 and SAMN08102610. Computer code used to analyze the data and produce the results in the paper are on GitHub at https://github.com/jbloomlab/Perth2009-DMS-Manuscript.

### HA numbering

Sites are in H3 numbering, with the signal peptide in negative numbers, HA1 in plain numbers, and HA2 denoted with "(HA2)". Sequential 1, 2, … numbering of the Perth/2009 HA can be converted to H3 numbering by subtracting 16 for the HA1 subunit, and subtracting 345 for the HA2 subunit.

### Creation of MDCK-SIAT1-TMPRSS2 cells

When growing influenza virus in cell culture, trypsin is normally added to cleave HA into its mature form. To obviate the need for trypsin, we engineered MDCK-SIAT1 cells to constitutively express the TMPRSS2 protease, which cleaves and activates HA in the human airways (30, 31). The human TMPRSS2 cDNA ORF was ordered from OriGene (NM_005656) and cloned into a pHAGE2 lentiviral vector under an EF1α-Int promoter followed by an IRES driving expression of mCherry to create plasmid pHAGE2-EF1aInt-TMPRSS2-IRES-mCherry-W. We used the lentiviral vector to transduce MDCK-SIAT1 cells, and sorted an intermediate mCherry-positive population by flow cytometry. We refer to the sorted bulk population as MDCK-SIAT1-TMPRSS2 cells. There is no selectable marker for the TMPRSS2; however, we maintain the cells at low passage number, and have seen no indication that they lose their ability to support the growth of viruses with H3 HAs in the absence of exogenous trypsin.

### Generation of HA codon mutant plasmid libraries

HA and NA genes for the Perth/2009 viral strain were cloned from recombinant virus obtained from BEI Resources (NR-41803) into the pHW2000 (81) influenza reverse-genetics plasmids to create pHW-Perth09-HA and pHW-Perth09-NA.

We initially created a virus with the HA and NA from Perth/2009 and internal genes from WSN/1933, and passaged it in cell culture to test its genetic stability. To generate this virus, we transfected a co-culture of 293T and MDCK-SIAT1-TMPRSS2 in D10 media (DMEM supplemented with 10% heat-inactivated FBS, 2 mM L-glutamine, 100 U of penicillin/mL, and 100 μ of streptomycin/mL) with equal amounts of pHW-Perth09-HA, pHW-Perth09-NA, the pHW18* series of plasmids (81) for all non-HA/NA viral genes, and pHAGE2-EF1aInt-TMPRSS2-IRES-mCherry-W. The next day we changed the media to influenza growth media (IGM, consisting of Opti-MEM supplemented with 0.01% heat-inactivated FBS, 0.3% BSA, 100 U of penicillin/mL, 100 *μ*g of streptomycin/mL, and 100 *μg* of calcium chloride/mL (no trypsin was added since there was TMPRSS2), and then collected the viral supernatant at 72 hours post-transfection. This viral supernatant was blind passaged in MDCK-SIAT1-TMPRSS2 a total of six additional times. We isolated viral RNA from these passaged viruses and sequenced the HA gene. The passaged HA had two mutations, G78D and T212I, which enhanced viral growth as shown in Figure S1. The HA with these two mutations was cloned into pHW2000 (81) and pICR2 (82) to create pHW-Perth09-HA-G78D-T212I and pICR2-Perth09-HA-G78D-T212I. For all subsequent experiments, we used viruses with the HA containing these two mutations to improve titers and viral genetic stability, and this is the HA that we refer to as Perth/2009 in the manuscript text. We used all non-HA genes (including NA) from WSN/1933 to help increase titers and reduce biosafety concerns.

The codon-mutant libraries were generated in the Perth/2009 HA-G78D-T212I background using the PCR-based approach described in (83) with the primer melting-temperature modifications described in (84), using two rounds of mutagenesis. The script to design the mutagenesis primers is at https://github.com/jbloomlab/ CodonTilingPrimers. We created three independent libraries, one for each biological replicate. The mutant variants were then cloned at high efficiency into the pICR2 (82) vector using digestion with BsmBI, ligation with T4 DNA ligase, and electroporation into ElectroMAX DH10B competent cells (Invitrogen 18290015). We obtained >6 million transformants for each replicate. We scraped the plates, expanded the cultures in liquid LB + ampicillin at 37°C for 3 h with shaking, and then maxiprepped. We randomly chose 31 clones to Sanger sequence to evaluate the mutation rate (Figure S2).

### Generation and passaging of mutant viruses

The mutant virus libraries were generated using the helper-virus approach described in (10) with several modifications, most notably the cell line used. Briefly, we transfected 5 x 10^5^ MDCK-SIAT1-TMPRSS2 cells in suspension with 937.5 ng each of four protein expression plasmids encoding the ribonucleoprotein complex (HDM-Nan95-PA, HDM-Nan95-PB1, HDM-Nan95-PB2, and HDM-Aichi68-NP) (75), and 1250 ng of one of the three pICR2-mutant-HA libraries (or the wild-type control) using Lipofectamine 3000 (ThermoFisher L3000008). We allowed the transfected cells to adhere in 6-well plates and four hours later changed the media to D10 media. Eighteen hours after transfection, we infected the cells with the WSN/1933 HA-deficient helper virus (10) by preparing an inoculum of 500 TCID50 per *μ*L of helper virus (as computed on HA-expressing cells) in IGM, aspirating the D10 media from the cells, and adding 2 mL of the helper-virus inoculum to each well. After three hours, we changed the media to fresh IGM. At 24 hours after helper-virus infection, we harvested the viral supernatants for each replicate, froze aliquots at −80°C, and titered in MDCK-SIAT1-TMPRSS2 cells. The titers were 92, 536, 536, and 734 TCID50 per *μ*L for the three library replicates and the wildtype control, respectively.

We passaged 9 x 10^5^ TCID50 of the transfection supernatants at an MOI of 0.0035 TCID50 per cell. To do this, we plated 4.6 x 10^6^ MDCK-SIAT1-TMPRSS2 cells per dish in 15-cm dishes in D10 media, and allowed the cells to grow for 24 hours, at which time they were at ~1.7 x 10^7^ cells per dish. We replaced the media in each dish with 25 mL of an inoculum of 2.5 TCID50 of virus per *μ*L in IGM. We collected viral supernatant for sequencing 48 hours post-infection.

### Barcoded subamplicon sequencing

To extract viral RNA from the three replicate HA virus libraries and the wildtype HA virus, we ultracentrifuged 24 mL of supernatant at 22,000 rpm for 1.5 h at 4^°^ C in a Beckman Coulter SW28 rotor, and extracted RNA using the Qiagen RNeasy Mini Kit by resuspending the viral pellet in 400 *μ*L of buffer RLT supplemented with *β*-mercaptoethanol, pipetting 30 times, transferring the liquid to a microcentrifuge tube, adding 600 *μ*L 70% ethanol, and proceeding according to the manufacturer’s instructions. The HA gene was reverse-transcribed with AccuScript Reverse Transcriptase (Agilent 200820) using primers P09-HA-For (5’-AGCAAAAGCAGGGGATAATTCTATTAATC-3’) and P09-HA-Rev (5’-AGTAGAAACAAGGGTGTTTTTAATTACTAATACAC-3’).

We generated the HA PCR amplicons for the three plasmid libraries, the three virus libraries, the wildtype plasmid control, and the wildtype virus control using KOD Hot Start Master Mix (EMD Millipore 71842) using the PCR reaction mixture and cycling conditions described in (83) and the P09-HA-For and P09-HA-Rev primers. We prepared the sequencing libraries using a barcoded-subamplicon strategy (24) to increase the accuracy from deep sequencing. The exact details of this approach are described in (10) (also see https://jbloomlab.github.io/dms_tools2/bcsubamp.html). The primers used to generate the subamplicons are in Dataset S2. We performed deep sequencing on a lane of an Illumina HiSeq 2500 using 2 x 250 bp paired-end reads in rapid-run mode.

### Analysis of deep sequencing data

We used the dms_tools2 software package (38) (https://github.com/jbloomlab/dms_tools2, version 2.2.0) to analyze the deep sequencing data. The algorithm used to estimate the site-specific amino-acid preferences from the deep sequencing counts is described in (38). The amino-acid preferences for each replicate and for the re-scaled, across-replicate average are provided in Dataset S3. Computer code that performs the entirety of this analysis and shows many detailed plots about read depth and other quality control metrics is available on GitHub at https://github.com/jbloomlab/Perth2009-DMS-Manuscript/blob/master/ analysis_code/analysis_notebook.ipynb.

### Phylogenetic model comparison and fitting of a stringency parameter

For the analysis in Table 1, we downloaded all full-length H3 HA sequences from the Influenza Virus Resource (85), and randomly subsampled two sequences per year. These sequences were aligned using MAFFT (86) and used to infer a phylogenetic tree using RAxML (87) with a GTRCAT model of nucleotide substitution. We then used phydms (34) (https://github.com/jbloomlab/phydms, version 2.2.1) to fit the substitution models listed in Table 1. The amino-acid preferences were re-scaled by the stringency parameter using the approach described in (34).

The phylogenetic tree of HA subtypes in Figure 7A was generated as described in (57).

### Quantification of relative solvent accessibility

We quantified the absolute solvent accessibility of each site of the H3 HA (PDB 4O5N; (40)) or the H1 HA (PDB 1RVX; (41)) structure using DSSP (88). We then normalized to a relative solvent accessibility using the absolute accessibilities in (89).

### Quantification of mutational effects

We calculated the effects of mutations from the amino-acid preferences that were estimated from the deep mutational scanning data. The effect of mutating site *r* from amino-acid *a_1_* to *a*_2_ was calculated as

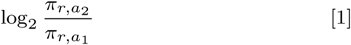

where π_*r,a1*_ and π _*r,a2*_ are the re-scaled preferences for amino acids *a*_1_ or a_2_ at site *r* as shown in Figure 2. The WSN/1933 H1 HA amino-acid preferences are the replicate-average values reported in (10), re-scaled by a stringency parameter of 2.05 (see https://github.com/jbloomlab/dms_tools2/blob/master/examples/Doud2016/analysis_notebook.ipynb).

### Inference of human H3N2 phylogenetic tree and calculation of maximum mutation frequencies

To generate the tree shown in Figure S4, we applied Nextstrain’saugur pipeline (90) (https://github.com/<u>nextstrain/augur;</u> commit 006896d) to publicly available H3N2 HA sequences from GISAID (91) (see Dataset S4), sampling six viruses per month over the time interval of January 1, 1968 to February 1, 2018. We aligned the resulting 2,189 HA sequences with MAFFT v7.310 (86) and constructed a maximum likelihood phylogeny from this alignment with RAxML 8.2.10 (87). Ancestral state reconstruction and branch length timing were performed with TreeTime (92). The phylogenetic tree is available as a JSON file on GitHub at https://github.com/jbloomlab/Perth2009-DMS-Manuscript/blob/master/ <u>analysis_code/data/flu_h3n2_ha_1968_2018_6v_tree.json.gz.</u> The tree was visualized using BALTIC (https://github.com/blab/baltic).

The frequency trajectory of each individual mutation on the phylogeny is estimated following Nextstrain’s augur pipeline and as first implemented in Nextflu (93). Herein, mutation frequency dynamics are modeled according to a Brownian motion diffusion process discretized to one-month intervals. Relative to a simple Brownian motion, the expectation includes an “inertia” term *ε* that adds velocity to the diffusion and the variance includes a term *x*(1 — *x)* to scale variance according to frequency following a Wright-Fisher population genetic process. This results in the following diffusion process with

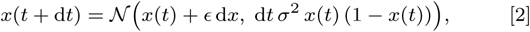

with ‘volatility’ parameter *σ*^2^. The term dx is the increment in the previous timestep, so that d*x* = *x(t) — x(t — dt).* We used *ε* = 0.7 and *σ^2^* = 0.05 to maximize fit to empirical trajectory behavior.

We also include an Bernoulli observation model for mutation presence / absence among sampled viruses at timestep *t*. This observation model follows

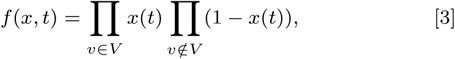

where *v* ∈ *V* represents the set of viruses that have the mutation and *v ∉ V* represents the set of viruses that do not have the mutation. Each frequency trajectory is estimated by simultaneously maximizing the likelihood of the process model and the likelihood of the observation model via adjusting frequency trajectory *x* = (x_1_,…,x_n_).

### Analysis of mutational shifts

To compare the Perth/2009 H3 and WSN/1933 H1 HA preferences, we first aligned the wildtype HA sequences using MAFFT (86). To quantify the shifts in preference for every alignable site while accounting for experimental noise, we used the approach described in (27) and used the RMSDcorrected values as our quantification of the extent of each shift.

For the plots shown in Figure 8B, any residues falling between Cys-52 and Cys-277 were defined as the head domain, and all other residues were defined as the stalk domain. We used the multiple sequence alignment of the HA subtype sequences from (57) to identify sites that are absolutely conserved across all subtypes, or in the different clades described in Figure 8.

## ACKNOWLEDGMENTS

We thank Sarah Hilton, Hugh Haddox, and Sidney Bell for helpful discussions about data analysis, and Richard Neher for sharing analysis code and providing helpful comments on the manuscript. We thank the Fred Hutch Genomics Core for performing the Illumina deep sequencing. This work was supported by grant R01 AI127893 from the NIAID of the NIH to JDB and TB and grant U19 AI117891 to TB. JML was supported in part by the Center for Inference and Dynamics of Infectious Diseases (CIDID), which is funded by grant U54GM111274 from the NIGMS of the NIH. The research of JDB is supported in part by a Faculty Scholar grant from the Howard Hughes Medical Institute and the Simons Foundation, and a Burroughs Wellcome Young Investigator in the Pathogenesis of Infectious Diseases grant. TB is a Pew Biomedical Scholar and is supported by NIH R35 GM119774.

## Supporting Information (SI)

**Figure S1.**
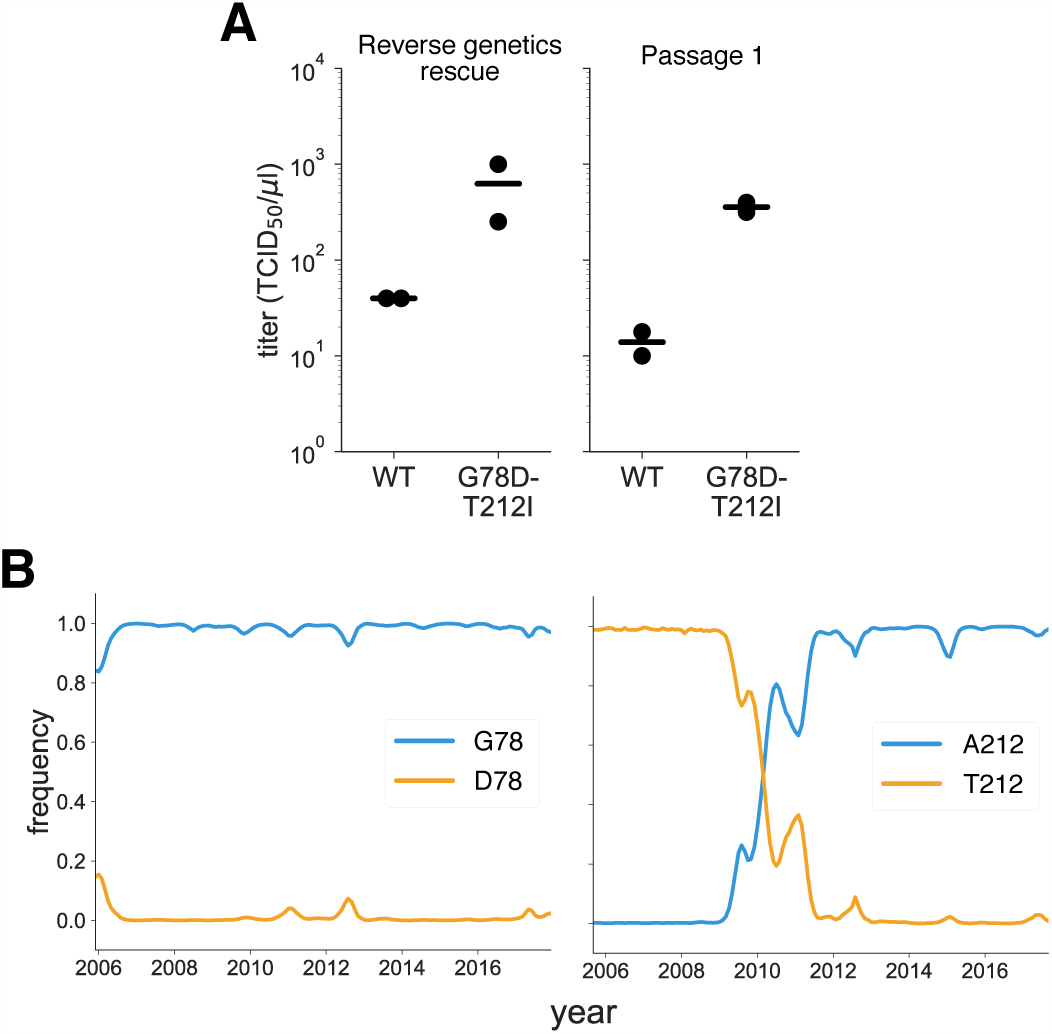
Characterization of the G78D-T212I Perth/2009 HA variant. (A) The G78D-T212I Perth/2009 HA variant supports better viral growth than the wildtype Perth/2009 HA. Viruses were generated in duplicate by reverse genetics with the Perth/2009 NA and WSN internal genes, and passaged once at MOI = 0.01 in MDCK-SIAT1-TMPRSS2 cells. The rescue and passage viral supernatants were collected at 72 hours post-transfection and 44 hours post-infection, respectively, and titered in MDCK-SIAT1-TMPRSS2 cells. The points mark each duplicate and the bar marks the mean. (B) The D78 variant remained at a low frequency in natural human H3N2 sequences over the past ~10 years. The A212variant rose to fixation in ~2011, replacing the T212 variant.

**Figure S2.**
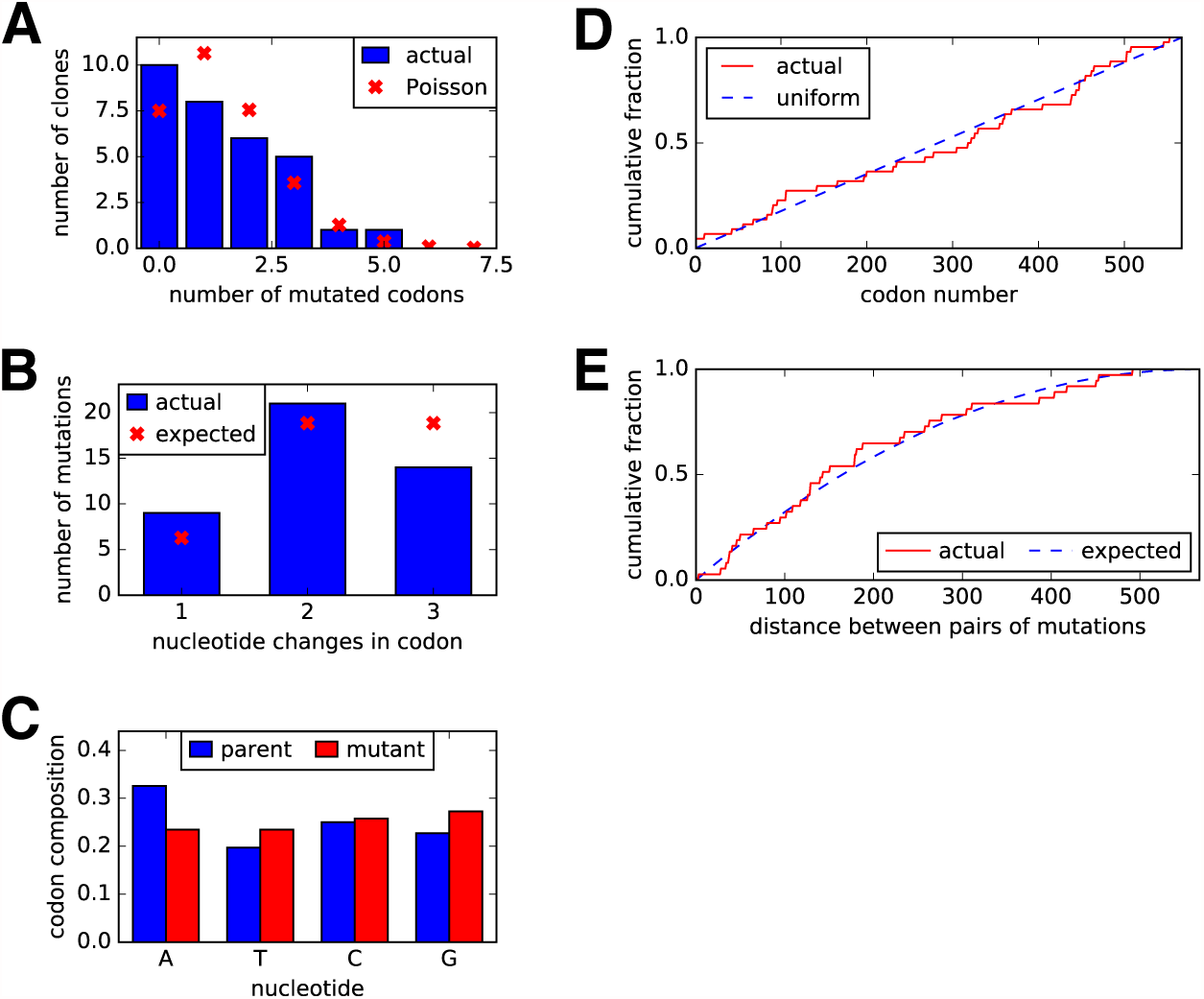
Sanger sequencing of 31 randomly chosen clones from the mutant plasmid libraries. (A) There were an average of ~1.4 codon mutations per clone across the three plasmid mutant libraries. (B) A mixture of one-, two-, and three-nucleotide mutations were present, with slightly fewer triple-nucleotide changes than expected. (C) Nucleotide frequencies were uniform in the codon mutations. (D) The mutations were distributed relatively evenly across the length of the HA coding sequence. (E) We calculated the pairwise distances between mutations for clones carrying more than one mutation. The cumulative distribution of these distances is shown in the red line. The blue line indicates the expected distribution if mutations in multiply mutated genes are randomly dispersed along the sequence.

**Figure S3.**
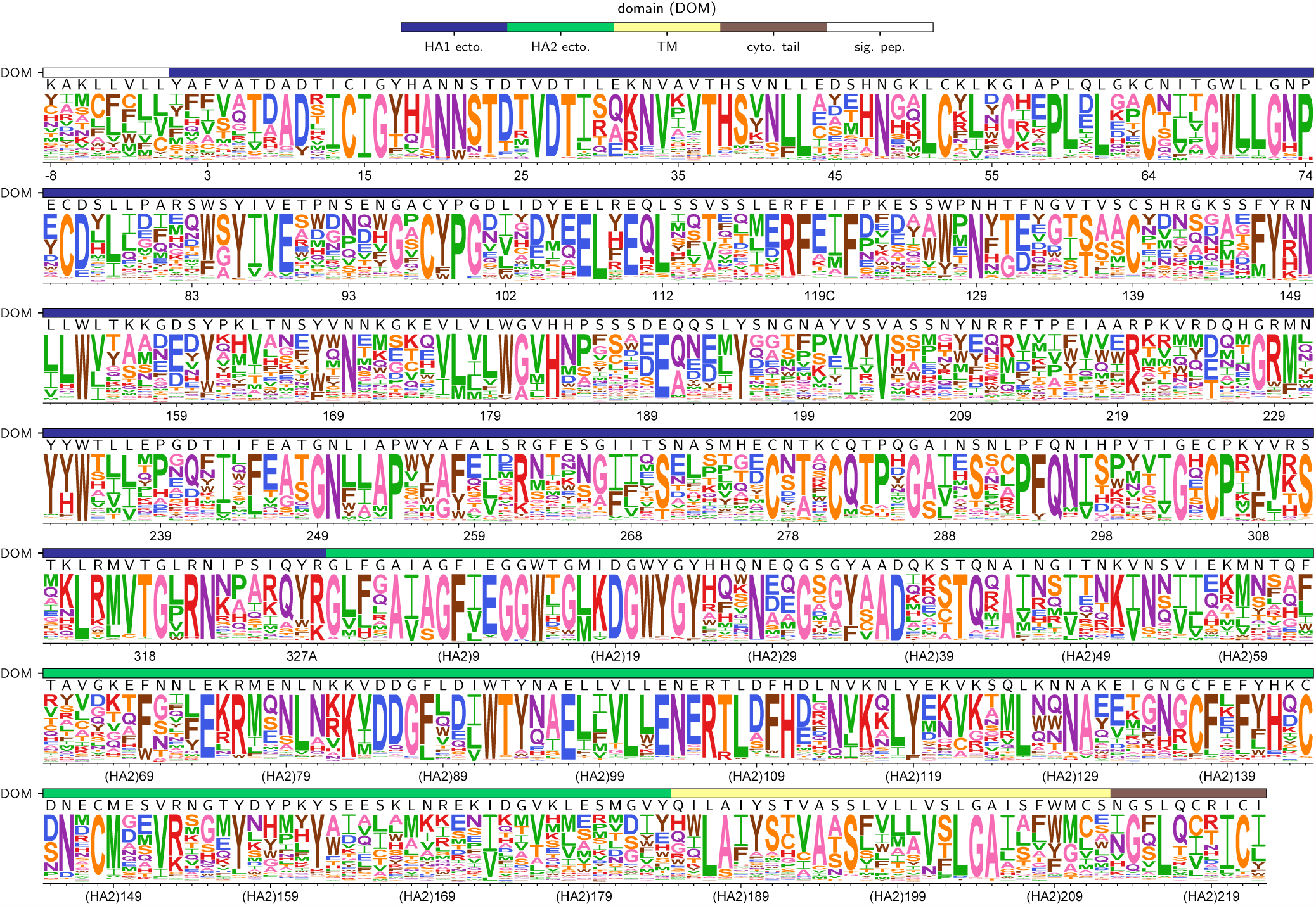
The site-specific amino-acid preferences of the WSN/1933 H1 HA as measured in (10). The amino-acid preferences from (10) after taking the average of the experimental replicates and re-scaling (34) by a stringency parameter of 2.05 (see https://github.com/jbloomlab/dms_tools2/blob/master/examples/ <u>Doud2016/analysis_notebook.ipynb)</u>. The sites are in H3 numbering. The overlays show the same information as in Figure 2 (domain and wildtype amino acid).

**Figure S4.**
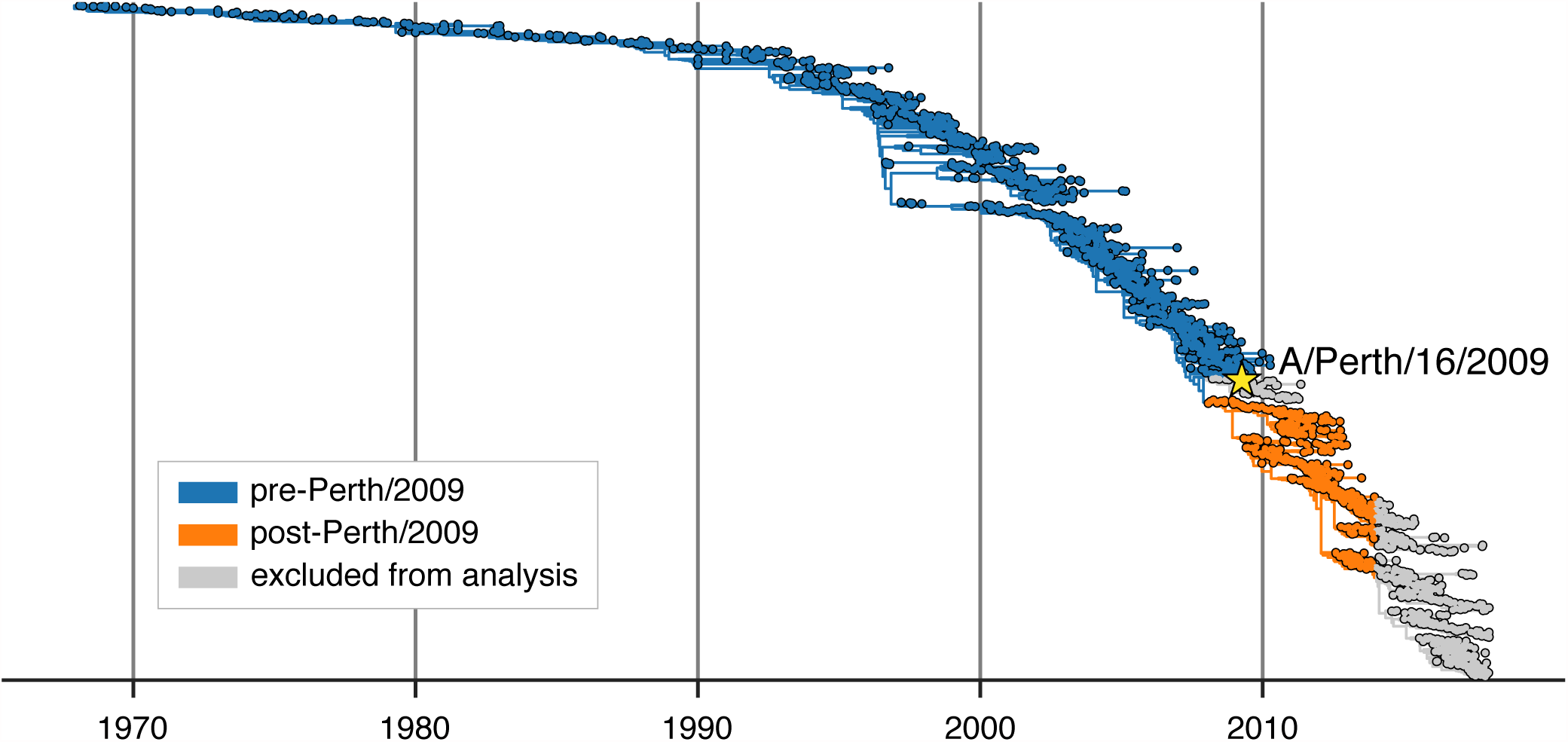
A phylogenetic tree of all HA sequences used in our analysis of mutation frequencies. HA sequences were sampled at a rate of six viruses per month from January 1, 1968 through February 1,2018. The Perth/2009 strain used in our experiments is indicated. The rest of the tree is partitioned into nodes that preceded the split of the Perth/2009 strain from the trunk of the tree (blue) and nodes that branched off the trunk after the clade containing Perth/2009 (orange). In Figure 5, these two partitions of the tree are analyzed separately. Nodes in the clade containing the Perth/2009 strain and nodes sampled in 2014 or after were excluded from our analyses. The Perth/2009 strain was excluded to avoid artifacts related to mutations that occurred on the branches leading to the HA sequence used in the experiment. The post-2014 nodes were excluded because the evolutionary fates of many sequences after this date are not yet full resolved.

**Figure S5.**
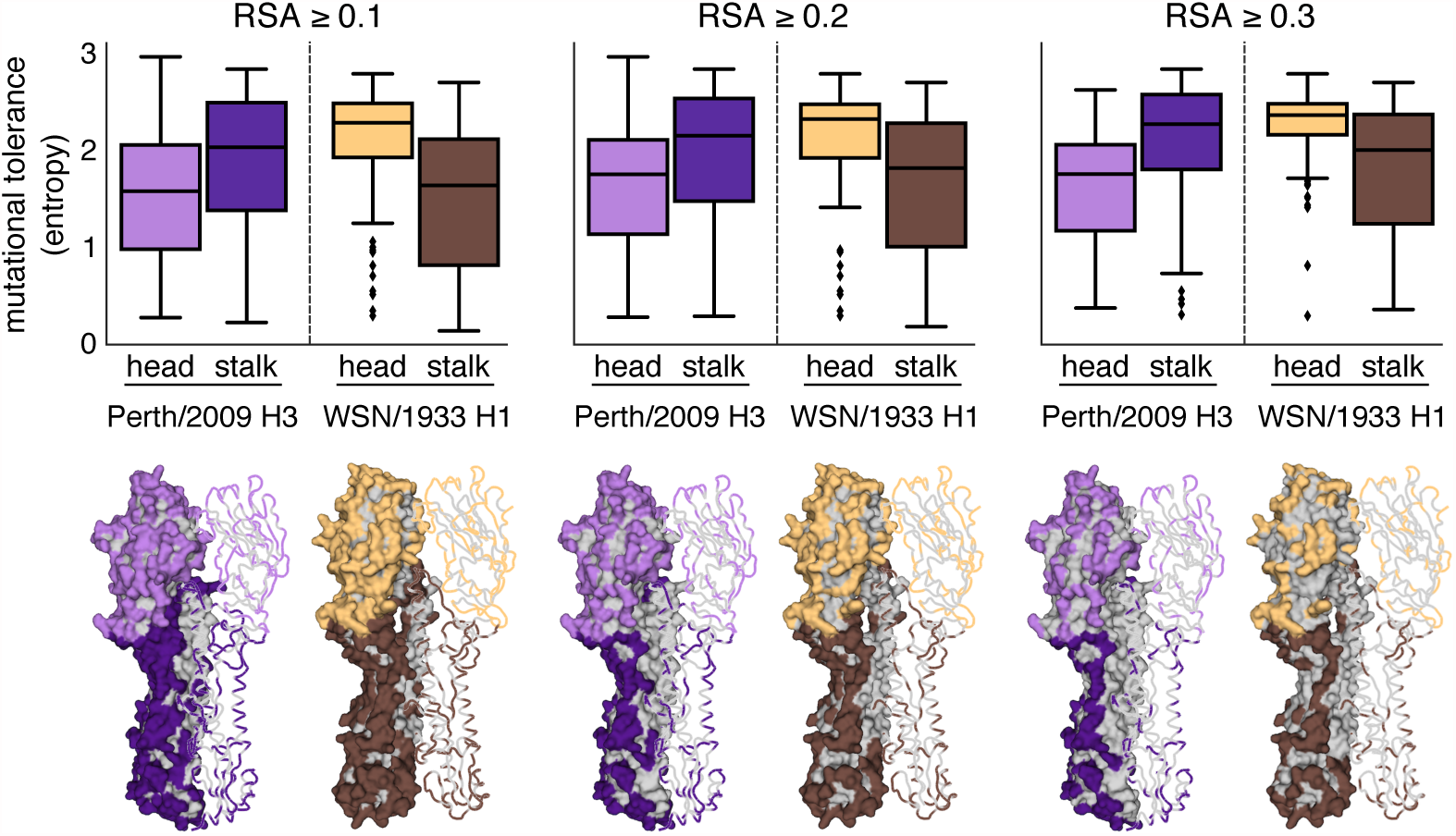
Mutational tolerances of the head and stalk domains at various relative solvent accessibility cutoffs. The mutational tolerances of the head and stalk domains show less disparity for the Perth/2009 H3 HA compared to those for the WSN/1933 H1 HA. We used relative solvent accessibility (RSA) cutoffs of 0.1, 0.2, and 0.3 to define solvent-exposed residues and plotted the mutational tolerances (Shannon entropy of re-scaled preferences) of these residues in the head and stalk domains for the Perth/2009 H3 HA (purple) and the WSN/1933 H1 HA (brown). Residues falling in between the two cysteines at sites 52 and 277 were defined as belonging to the head domain, while all other residues were defined as the stalk domain. The HA structures color the residues that are defined as solvent exposed at a given RSA cutoff. One monomer is shown in surface representation and another monomer shown in ribbon representation. Residues in lighter shades of purple or brown are in the head domain, while residues in darker shades are in the stalk domain. Note that the mutational tolerance values are not comparable between the two HAs.

**Figure S6.**
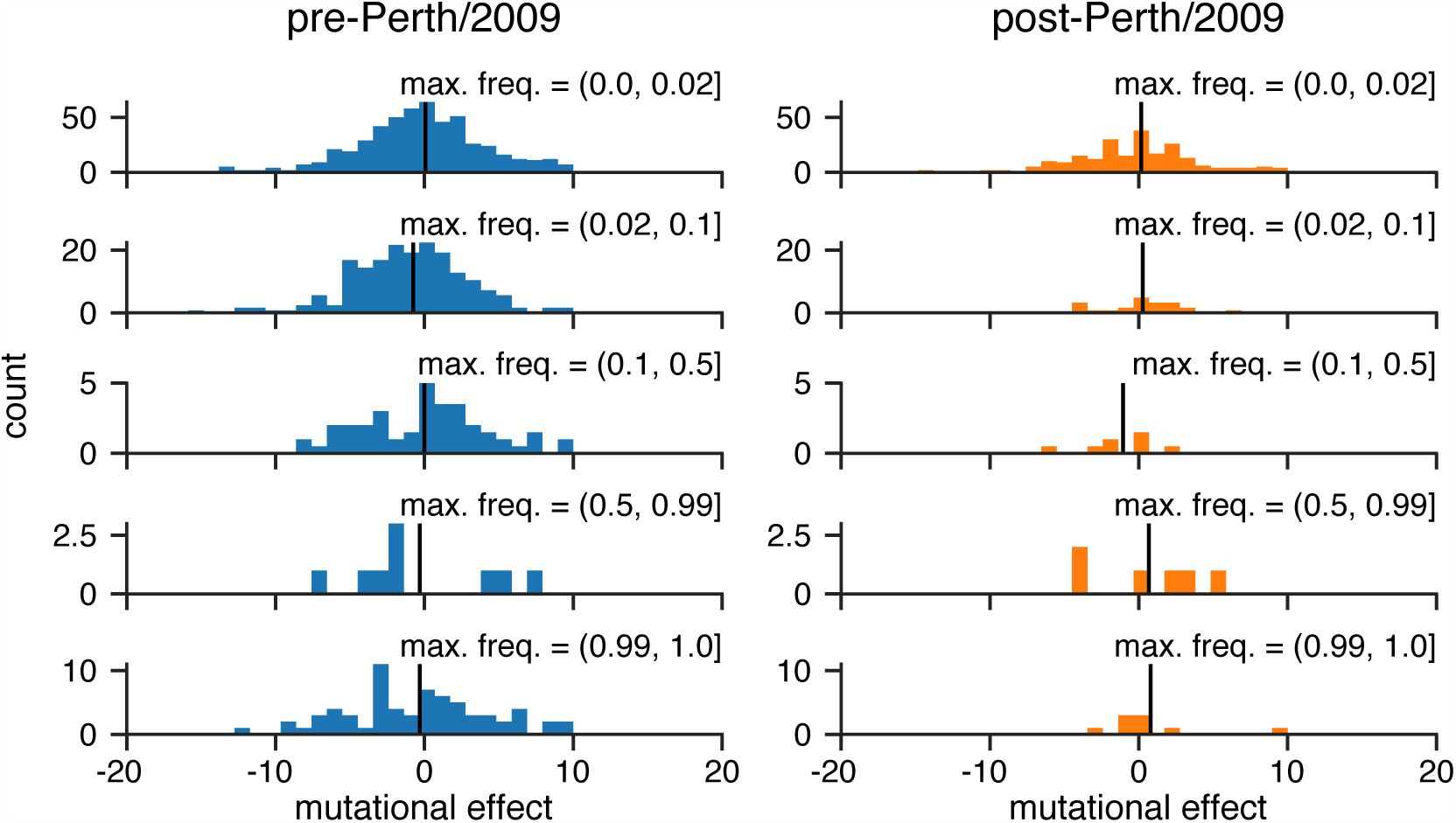
The distribution of mutational effects measured in H1 HA among H3N2 mutations binned by the maximum frequency that they reach. This figure repeats the analysis of the H3N2 mutation frequencies in Figure 5B, but uses the deep mutational scanning data for an H1 HA as measured in (10).

**Dataset S1.** Genbank file giving the full sequence of the bidirectional reverse-genetics plasmid pHW-Perth09-HA-G78D-T212I, which encodes the wildtype HA sequence used in this study.

**Dataset S2.** Excel file providing the primers used to generate the barcoded subamplicons for Perth/2009 HA deep sequencing.

**Dataset S3.** Excel file giving the amino-acid preferences in sequential 1, 2,… numbering of the Perth/2009 HA. The unscaled preferences for replicates 1, 2, 3-1, and 3-2 are each in a separate tab of the file. Additional tabs give the across-replicate averaged and re-scaled amino-acid preferences in sequential numbering and in H3 numbering as shown in Figure 2. There are also tabs that give the conversion from sequential to H3 numbering. Each tab can simply be exported to CSV for computational analyses.

**Dataset S4.** Excel file providing acknowledgments and accessions for sequences downloaded from GISAID.

